## Supplementary figures for "Sex-biased regulatory changes in the placenta of native highlanders contribute to adaptive fetal development"

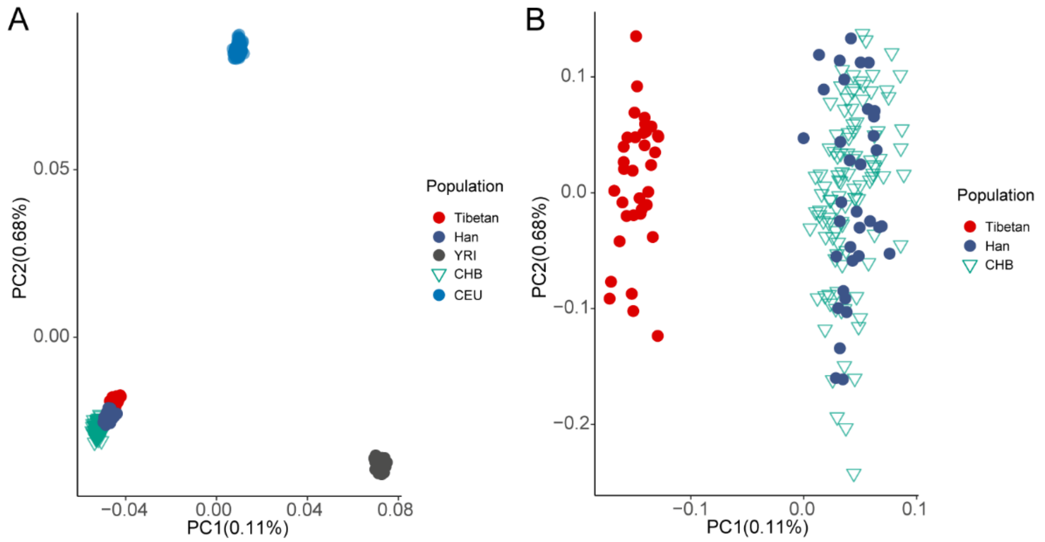


**Figure S1. The PCA plot of the 69 individuals in this study. (A)** The PCA plot of the 69 individuals (35 Tibetans and 34 Han migrants) based on the genome-wide variants, indicating their East Asian ancestry. **(B)** The zoom-in PCA plot displays the genetic divergence between Tibetans and Han, and no admixture is seen in the studied 69 individuals.

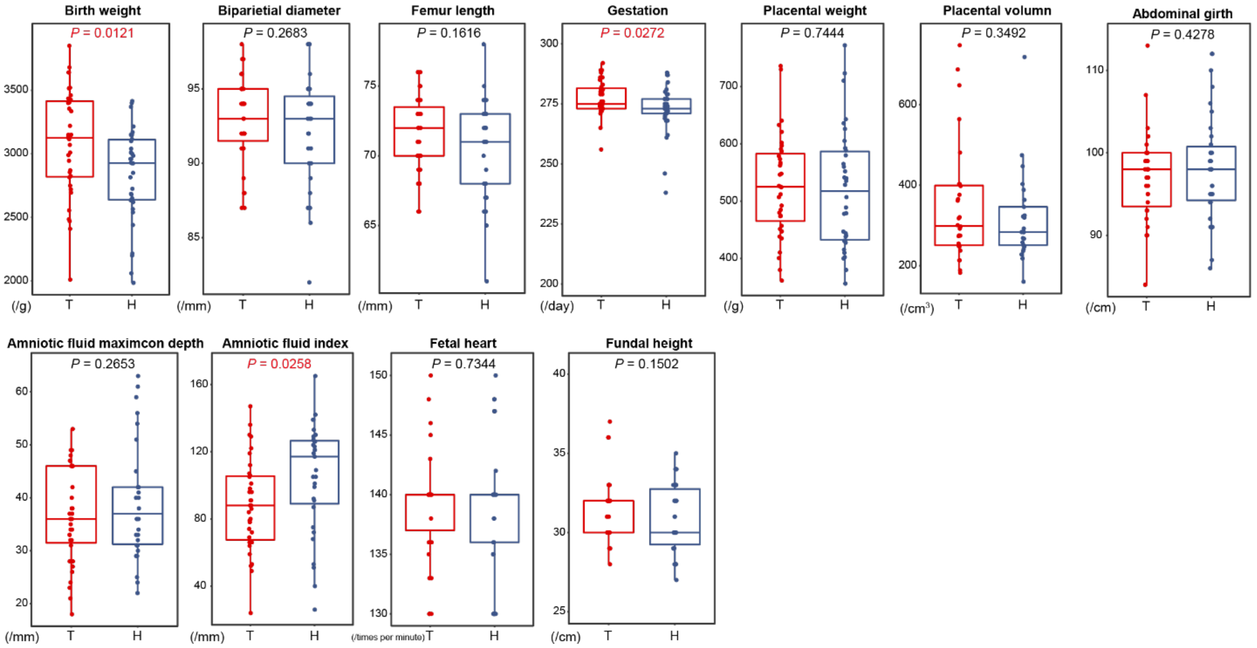


**Figure S2. Comparison of 11 reproductive traits between 35 Tibetans and 34 Han immigrants.** Univariate comparisons of the average value of each trait cross population were made by using the ANCOVA test in R *aov* function with fetal sex and maternal age as covariates.

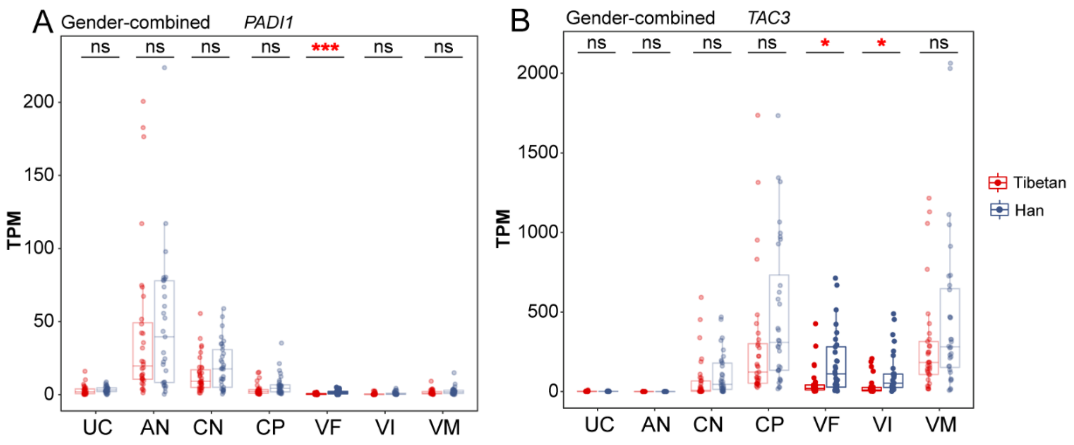


**Figure S3. The top DEGs of the VI and VF layers between Tibetans and Han.** Comparison of the expression level of *PADI1* **(A)** and *TAC3* **(B)** in the seven layers of placenta between Tibetans and Han. Adjusted *p*-value (*p*): *-$p < 0.05$; ***-$p < 0.001$; ns: not significant.

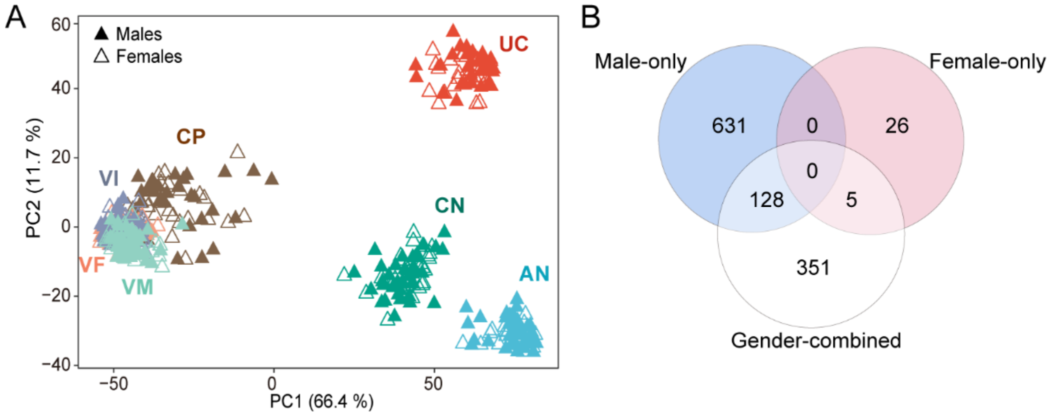


**Figure S4. The sex-biased gene expression in the placenta. (A)** The PCA map PCA shows the clustering pattern of tissue layers with fetal sex. **(B)** The overlapped DEGs between male-only group (blue), the female-only group (pink) and the gender-combined group (white).

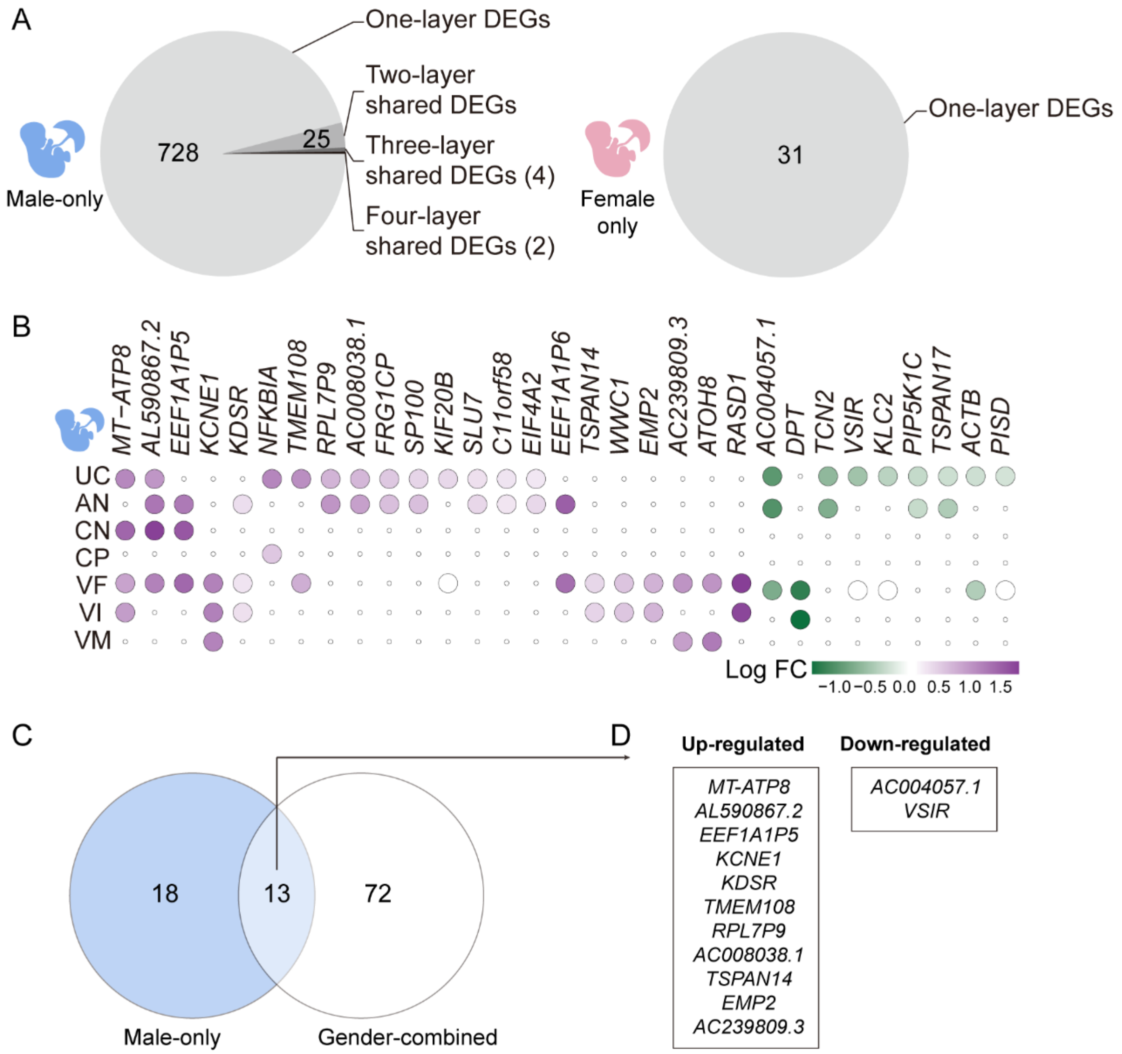


**Figure S5. The layer-shared DEGs in the placentas of males and females. (A)** The pie chart of the shared DEGs among two and more placental layers in males (left) and females (right). **(B)** The heat map of the 31 shared DEGs among two and more placental layers. Purple: up-regulated in Tibetan; green: down-regulated in Tibetans. **(C)** The overlap of layer-shared DEGs of the male-infant placentas and the gender-combined placentas. **(D)** The list of the 13 overlapped DEGs in **(C)**.

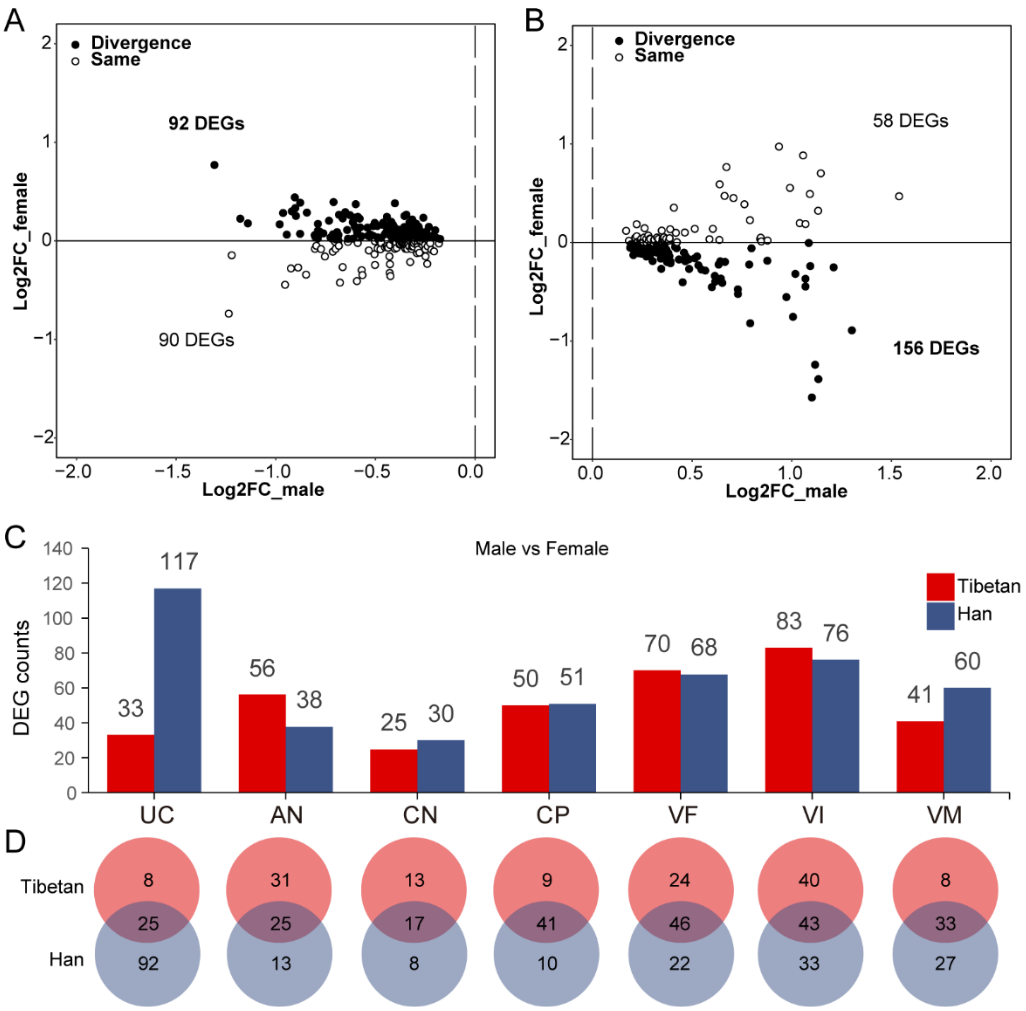


**Figure S6. The expression divergence between males and females. (A)** The differential direction of the down-regulated DEGs in the male UC layer in Tibetans and Han female infants. Solid circle: the differential expression direction of the male-DEGs are divergent in male and female placentas between Tibetans and Han; hollow circle: the differential expression direction of the male-DEGs are the same in male and female placentas between Tibetan and Han. **(B)** The differential direction of the up-regulated DEGs in the male UC layer in Tibetan and Han female infants. **(C)** Bar plot of DEGs (male-infant placenta vs female-infant placenta) of the seven layers of placenta in Tibetans and Han migrants, respectively. **(D)** The overlapped DEGs (male-infant placenta vs female-infant placenta) between Tibetans and Han migrants in the seven layers of placenta.

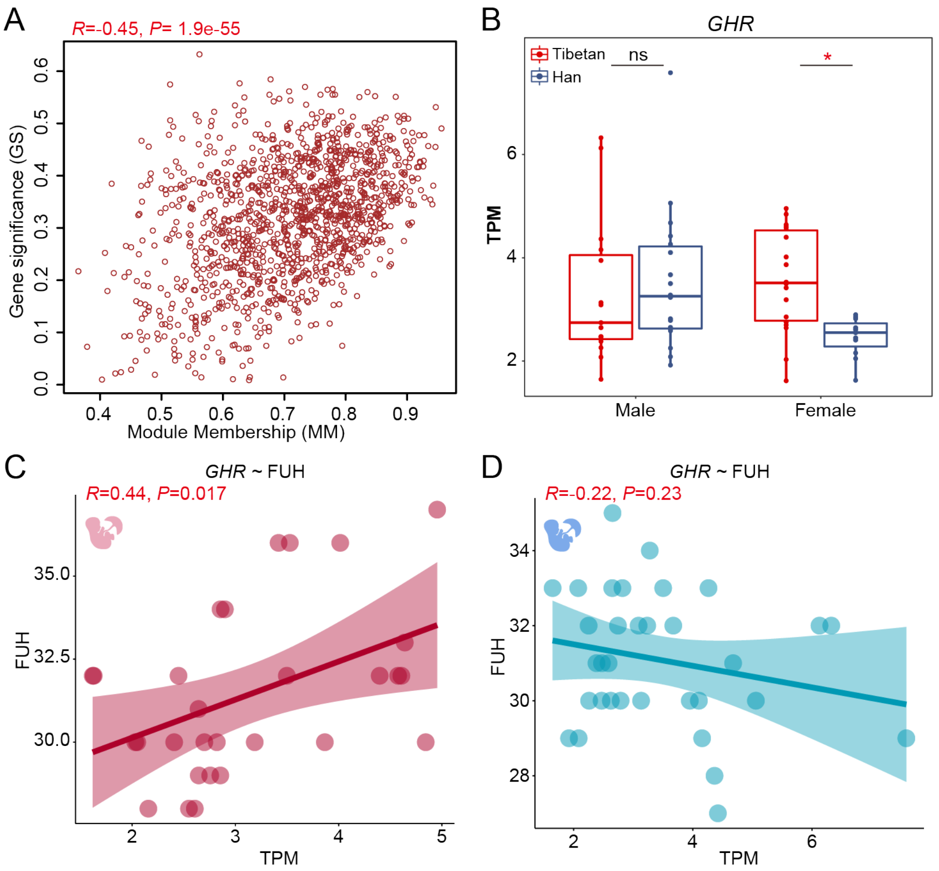


**Figure S7. The sex-biased correlation between gene expression and traits of the male-infant and female-infant placentas. (A)**The correlation between gene expression and BW in the male Module 8. Gene Significance (GS): the absolute value of the correlation between the gene and the trait; Module membership (MM): the correlation of the module eigengene and the gene expression profile. **(B)** Boxplot of the *GHR* gene in the male and female VF layers. **(C)** Correlation of *GHR* and FUH in the VF layer of the females. **(D)** Correlation of *GHR* and FUH in the VF layer of males.

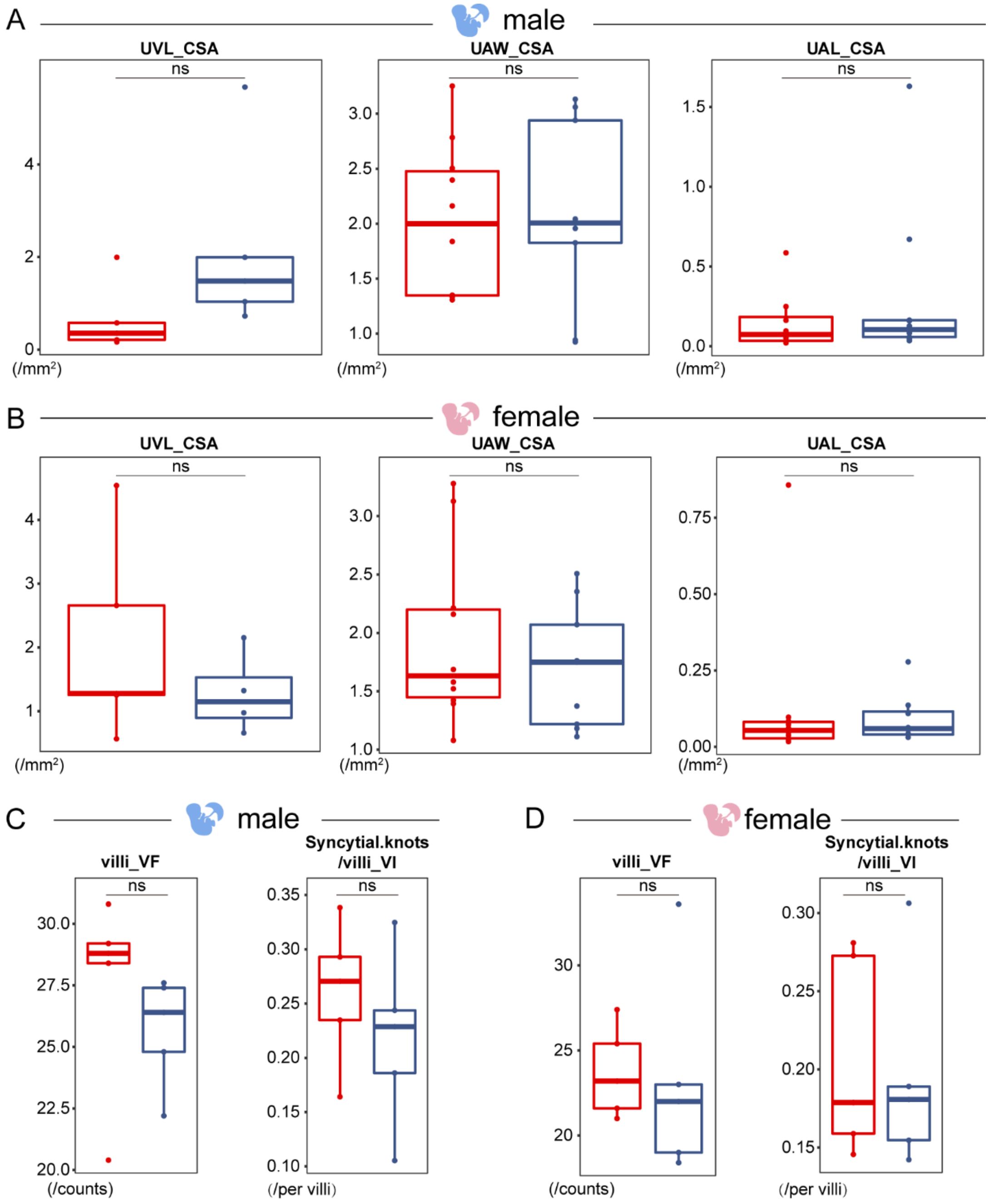


**Figure S8. Histological outcome of the sex-biased structure divergence in the placenta. (A)** Three parameters of UC in the male placentas. **(B)** Three parameters of UC in the female placentas. **(C)** Villi of the VF layer and syncytial knots per villi of the VI layer in the male placentas. **(D)** Villi of the VF layer and syncytial knots per villi of the VI layer in the female placentas. ns: not significant.

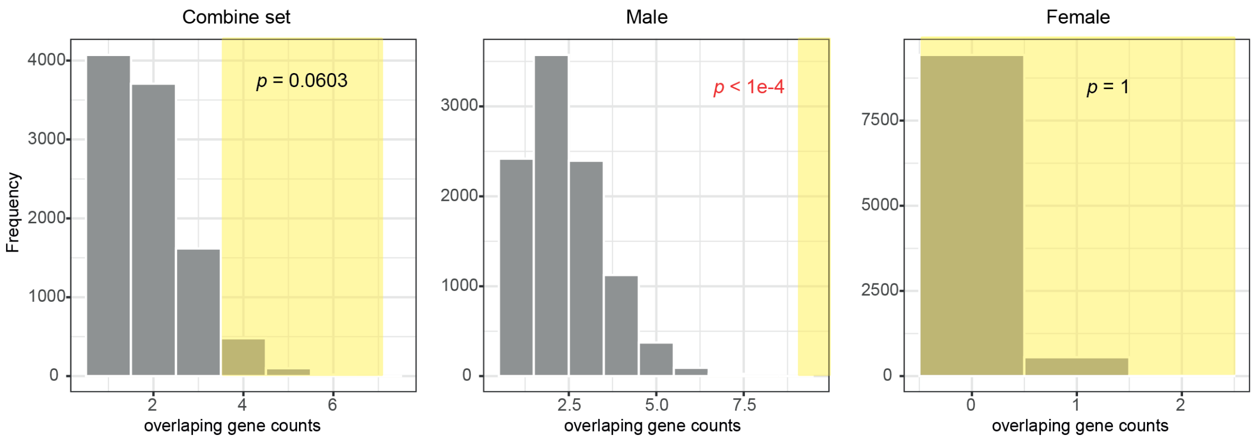


**Figure S9.** The distribution of the overlapped genes between the identified DEGs and the 192 randomly selected genes based on 10,000 permutations. The highlighted parts indicate the counts in the replicates with equal or more overlapping genes than the observed (≥4 for the combined set; ≥9 for the male-only set; ≥0 for the female-only set)

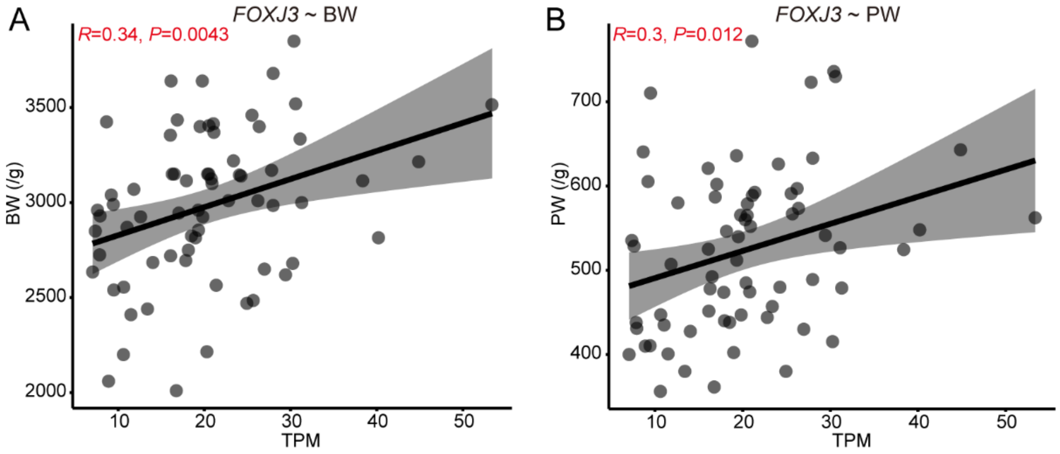


**Figure S10. The correlation between the expression of *FOXJ3* and BW/PW in the CN layer of placenta. (A)** The correlation between *FOXJ3* gene expression and BW. **(B)** The correlation between *FOXJ3* gene expression and PW.

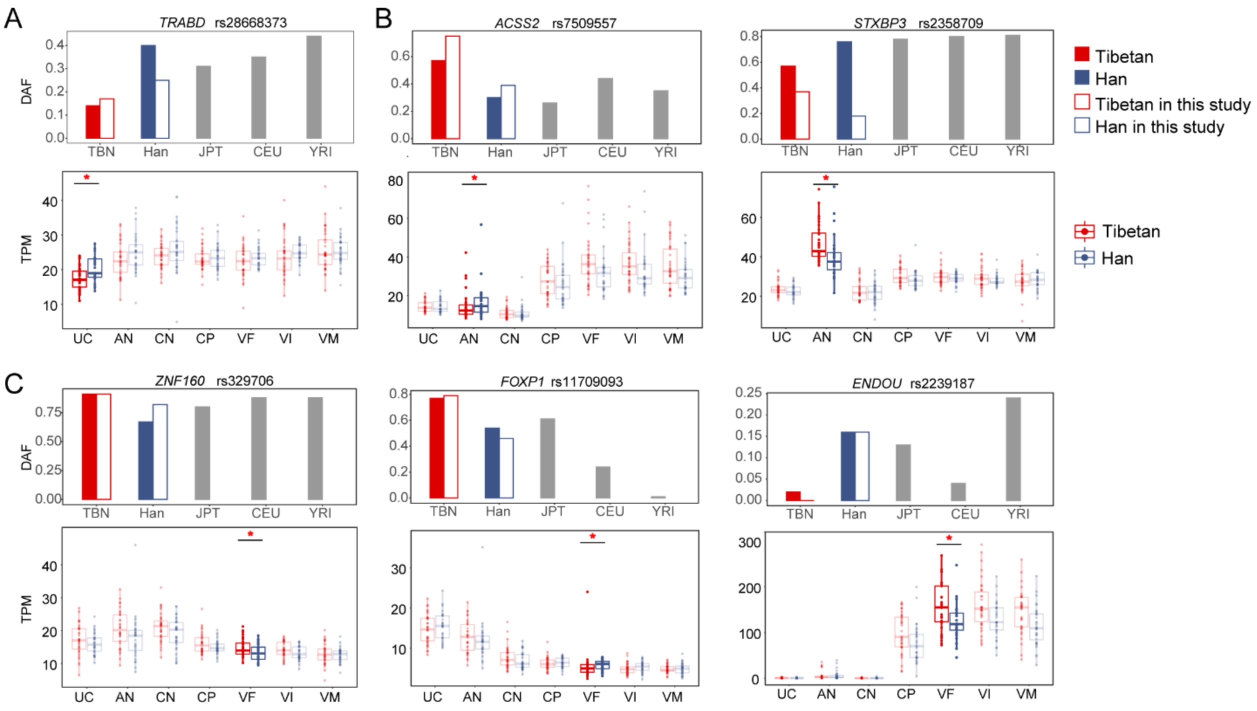


**Figure S11. The placental DEGs of the AN and VF layers with signals of positive selection in Tibetan population.** (A) *TRABD* (gender-separated analysis) of the UC layer with signals of positive selection in Tibetans. The upper panels show the allele frequencies of variant with the top signals of positive selection within the gene. Besides of Tibetans and Han Chinese, the frequencies of other three reference populations are also presented, including Japanese (JPT), Europeans (CEU) and Africans (YRI). The solid and hollow bars in red denote the allele frequencies in the published 1001 Tibetan individuals and 35 Tibetan individuals, respectively, and the solid and hollow bars in blue denote the allele frequencies in the published 103 Han individuals and 34 Han individuals, respectively. The bottom panels show the comparison of the expression levels between Tibetans and Han in the seven layers of placenta. Only the significant between-population differences are indicated. adjusted *p*-value: *, $p$-value < 0.05; **, $p$-value < 0.01; ***, $p$-value < 0.001. **(B)** The two DEGs (gender-separated analysis) of the AN layer with signals of positive selection in Tibetans. **(C)** The three DEGs (gender-separated) of the VF layer with signals of positive selection in Tibetans.
